# SpliceVarMech: A causal generative framework for mechanistic interpretation of non-canonical splicing variants

**DOI:** 10.64898/2026.11.19.767866

**Authors:** Moustafa Zein, Aboul Ella Hassanien

## Abstract

Non-canonical splicing variants occur beyond the conserved dinucleotides flanking splice junctions and are difficult to interpret because disruption scores alone do not explain splice outcomes or quantify uncertainty. We have developed SpliceVarMech, a causal generative framework in which a dual-stream encoder compares the wild-type and mutant sequences and conditional discrete sequence diffusion generates candidate splice outcomes. We then apply Bayesian causal inference to estimate posterior disruption probabilities with 95% credible intervals. The framework reports the predicted splice mechanism together with sequence attribution maps and an exploratory mapping to clinical evidence labels. On a gold-standard cohort of experimentally validated variants from male infertility genes, Bayesian leave-one-out cross-validation on 31 variants yielded an area under the receiver operating characteristic curve of 1.000. We benchmarked 16 individual splice prediction tools on the same cohort, where the highest-performing tool, SpliceAI, achieved 0.619 on its 30 scored variants. Ablation experiments showed that removing individual components reduced discrimination, and a separate evaluation on 54 variants supported the contribution of fine-tuning. In a cross-domain assessment, fine-tuning on BRCA1 saturation genome editing data increased discrimination from 0.722 to 0.840 on 203 held-out variants without architectural modification. We assessed calibration in-sample, and held-out validation remains necessary. These results support further evaluation of paired-sequence generative modeling and Bayesian inference for mechanistic interpretation of splicing variants across disease contexts.

## INTRODUCTION

RNA splicing is a fundamental step in gene expression, and variants that disrupt this process are increasingly recognized as a major cause of Mendelian disease.[1–3] Canonical splice-site variants at the conserved GT/AG dinucleotides are reliably identified by clinical pipelines. Non-canonical splicing variants (NCSVs) occur beyond the *±*2 positions flanking splice junctions and remain largely unresolved.[4, 5] These variants can disrupt auxiliary splicing signals, including exonic splicing enhancers (ESEs),[6] intronic splicing enhancers (ISEs), branch points, and polypyrimidine tracts.[3] Despite advances in splice prediction,[7, 8] interpreting their transcript-level effects remains challenging.[9] Splice-altering intronic variants have been documented across multiple disease contexts, from beta-sarcoglycanopathy[10] to VPS13D-related disorder,[11] underscoring the clinical breadth of this problem.

The clinical impact of unresolved NCSVs is particularly acute in male infertility, which affects approximately 7% of men worldwide.[12, 13] The testis has a diverse transcriptome, with extensive tissue-specific alternative splicing regulated by RNA-binding proteins.[1, 14] Transcriptional regulators such as NKAPL also contribute to gene regulation during meiosis exit.[15] A systematic review of male infertility whole-exome and whole-genome sequencing studies reported an average VUS burden of 47% among studies applying ACMG classification; 34% of the identified genes had undergone functional validation.[16] A review covering male and female infertility documented 73 functionally validated splicing variants across 54 genes, including 27 non-canonical variants potentially overlooked because they were considered likely benign or of uncertain significance.[17] Variant interpretation also presents challenges in pediatric hearing loss,[18] while studies of X-linked Alport syndrome[19] and Fabry disease[5] illustrate the importance of investigating splice-altering variants. Reclassification of such variants using computational evidence requires calibrated predictions that meet ACMG/AMP standards.[20, 21]

Existing splice prediction tools provide complementary information for interpreting NCSVs. SpliceAI was trained on GENCODE-annotated pre-mRNA sequences to predict splice sites, with additional splice-junction annotations from GTEx.[7] The source infertility study nevertheless identified experimentally validated variants that were difficult to interpret using available predictors.[4] Its 14 negative controls had predicted splice disruption but normal RT-PCR findings, with SpliceAI scores reaching as high as 0.84 for benign variants. Tissue-aware tools such as Pangolin[22] and SpliceTransformer[23] predict tissue-specific splicing, MMSplice models several splice outcomes,[8] and SQUIRLS provides interpretable variant prioritization.[24] Genomic foundation models such as DNABERT-2,[25] Nucleotide Transformer,[26] and Evo[27] further expand sequence-based modeling. These advances motivate frameworks that bring predicted transcript outcomes, explicit biological pathways and posterior uncertainty into a common interpretation workflow.

Here we present SpliceVarMech, a framework integrating mechanistic prediction of splice outcomes with Bayesian inference. Our approach combines discrete sequence diffusion models (D3PM)[28, 29] with a structural causal model[30, 31] to generate candidate mRNA outcomes and estimate disruption probabilities with credible intervals. The sequence-modeling component builds on advances in generative biological sequence modeling,[32, 33] while posterior inference uses the No-U-Turn Sampler (NUTS).[34] An exploratory ACMG evidence mapping accompanies the predictions.[20, 21] We evaluate the framework on experimentally validated NCSVs from male infertility genes, compare it with the evaluated baseline tools, and assess adaptation to another disease domain.

## RESULTS

### SpliceVarMech architecture overview

Figure 1 provides a schematic overview of the paired sequence inputs, model architecture and prediction outputs of SpliceVarMech. The framework comprises three integrated modules. Module 1 is a biological diffusion model (*∼*12.1M parameters) built on the D3PM architecture[28] with an absorbing-state noise schedule. Protein diffusion methods include continuous structure models[35, 36] and discrete amino-acid sequence models such as EvoDiff.[37] SpliceVarMech operates directly on the discrete nucleotide alphabet to model splice outcomes. The model processes paired wild-type (WT) and mutant (MUT) pre-mRNA contexts through a dual-stream encoder with cross-attention, in which MUT queries attend to WT keys and values, enabling explicit detection of variant-induced changes in splice signal recognition. This design is motivated by the dependence of splicing on sequence-specific recognition of splice sites and regulatory elements.[3] Paired inputs allow the encoder to represent changes associated with a variant.

**Figure 1:**
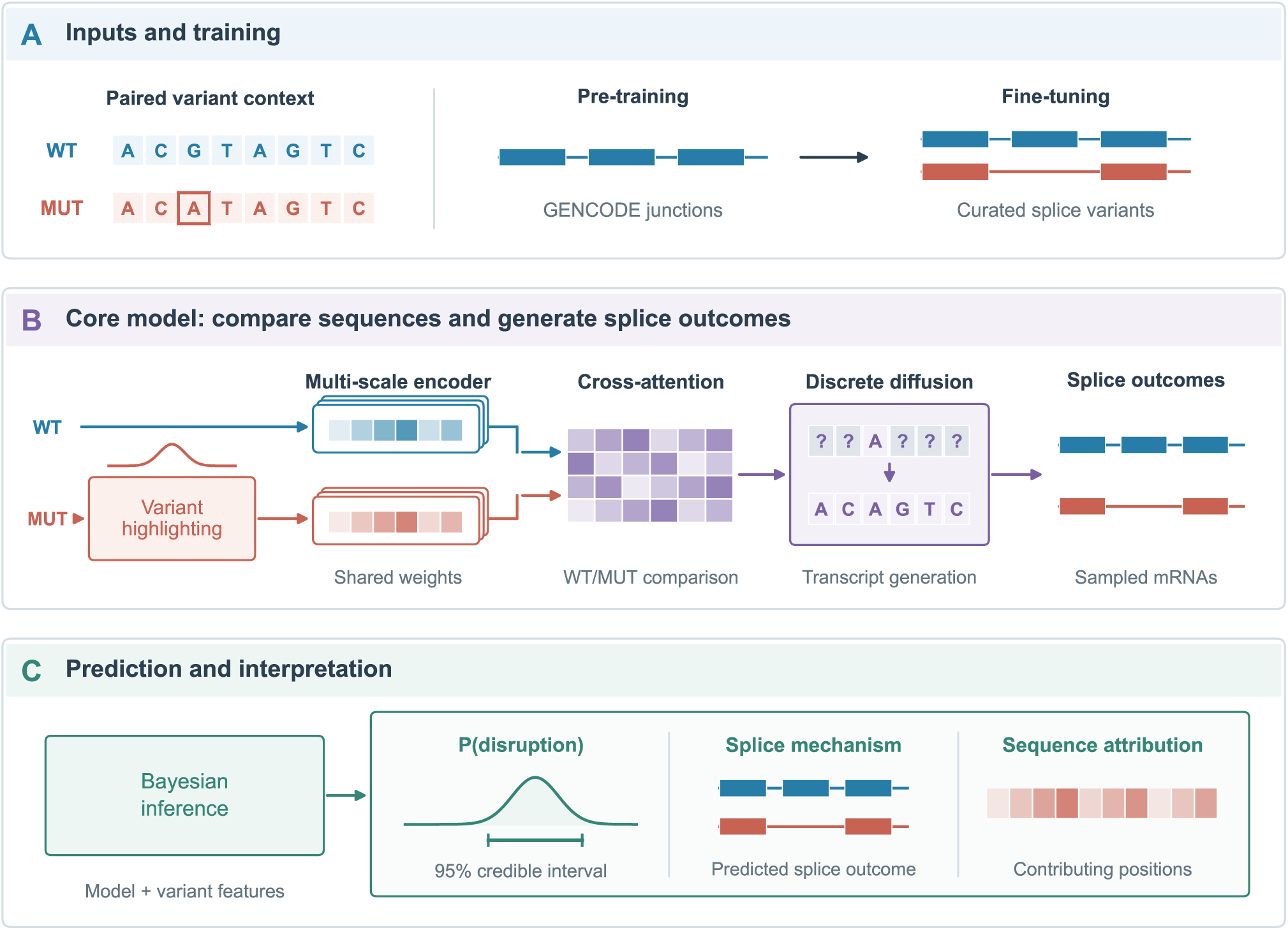
SpliceVarMech architecture. (A) Paired wild-type (WT) and mutant (MUT) sequence contexts provide the model inputs. Training comprises pre-training on reference splice junctions followed by fine-tuning on variant-containing contexts. (B) Variant highlighting, a shared multi-scale encoder and cross-attention compare WT and MUT sequences. Conditional discrete diffusion generates candidate splice outcomes from these representations. (C) Bayesian inference combines model-derived and splice features to estimate disruption probability with a 95% credible interval. Predicted splice mechanisms and sequence-attribution maps accompany the probability estimate. Sequence, mechanism and attribution illustrations are schematic. See also Table S4.

Multi-scale convolutional feature extraction uses kernels of 5, 15, and 51 nucleotides to capture local and extended sequence context. These architectural choices are motivated by the contributions of splice-site sequences,[38] exonic enhancer and silencer motifs,[6, 39, 40] and intronic regulatory elements.[3] The kernel widths are modeling choices rather than experimentally established boundaries for individual splicing factors. A contrastive learning objective enforces that disruptive variants produce distant WT/MUT representations while benign variants remain close. The D3PM reverse sampling process then generates predicted mRNA sequences, from which splice mechanism (exon skipping, intron retention, or cryptic site activation) and aberrant fraction are determined.

Module 2 is a Bayesian causal inference engine built on a structural causal model (SCM) representing relationships involved in splice site recognition.[30, 31] Bayesian network modeling has also been used for variant prioritization in Mendelian randomization.[41] Here, the SCM specifies pathways for variant-associated changes in splice strength (*V → S*), ESE/ESS balance (*V → E*), and ISE/ISS impact (*V → I*). MCMC inference via NUTS[34] produces posterior disruption probabilities with 95% credible intervals, expressing uncertainty conditional on the model and its inputs.

Module 3 generates explainable clinical reports through sequence attribution maps, causal path analysis, and ACMG criteria mapping,[20] producing a mechanism explanation, confidence grade, and reclassification recommendation for each variant.

### Two-stage training strategy

The model follows the two-stage training strategy outlined in Figure 1A. In Stage 1 (pre-training), the diffusion model learns to recognize canonical splice signals. These include GT/AG boundaries, ESE motifs,[6, 40] branch points, and polypyrimidine tracts. The model learns from exon-intron-exon splice junctions extracted from GENCODE v44 with a configured extraction limit of 252,835,[42] using tissue conditioning from GTEx v8 expression data across 10 tissue categories.[43] This pre-training stage teaches the model the “grammar” of normal splicing. It learns which sequence features the spliceosome requires for correct intron removal and how these requirements vary across tissues (Table S5).

In Stage 2 (fine-tuning), the model is adapted to variant effects using the 40 positive NCSVs and 14 negative controls in the source gold standard,[4] subject to sequence-context availability. The current diffusion-training loader also samples MFASS variants with minigene sequences (up to 200 disruptive and 200 benign variants, seed 42)[44] and admits up to 500 gnomAD v4.1 common intronic variants (AF*>*1%) with usable reference contexts.[45] These are sampling limits, not recorded final training counts. The model employs exponential moving average (EMA) weight smoothing,[46] self-conditioning,[47] and early stopping with a 15% validation holdout. The fine-tuning run reached the configured 30 epochs.

### Leave-one-out cross-validation on experimentally validated variants

We evaluated the Bayesian stage of SpliceVarMech using leave-one-out cross-validation (LOO-CV) on the 31-variant score-matched subset (23 pathogenic, 8 benign) of the full 54-variant gold standard (40 positives, 14 negatives).[4] For each held-out variant, the Bayesian model was retrained on the remaining 30 variants with MCMC inference (2,000 draws per chain, 2 chains, target acceptance 0.95). All 31 folds produced predictions, as summarized in Table S3. The saved fold-completion flags indicate successful model construction, sampling and prediction, but do not provide fold-wise *R*^^^ or effective sample size diagnostics.

SpliceVarMech achieved an AUROC of 1.000 and AUPRC of 1.000 (Figure 2C; Table 1). At the optimal threshold (*t* = 0.41, selected by maximizing Youden’s J statistic[48]), the framework correctly classified all 23 pathogenic variants and all 8 benign variants, yielding a sensitivity of 1.000, specificity of 1.000, balanced accuracy of 1.000, and MCC of 1.000 (Figure 2B). Perfect AUPRC (1.000) despite the class imbalance (74% positive prevalence) indicates that the framework maintains high precision at all recall operating points.

**Figure 2:**
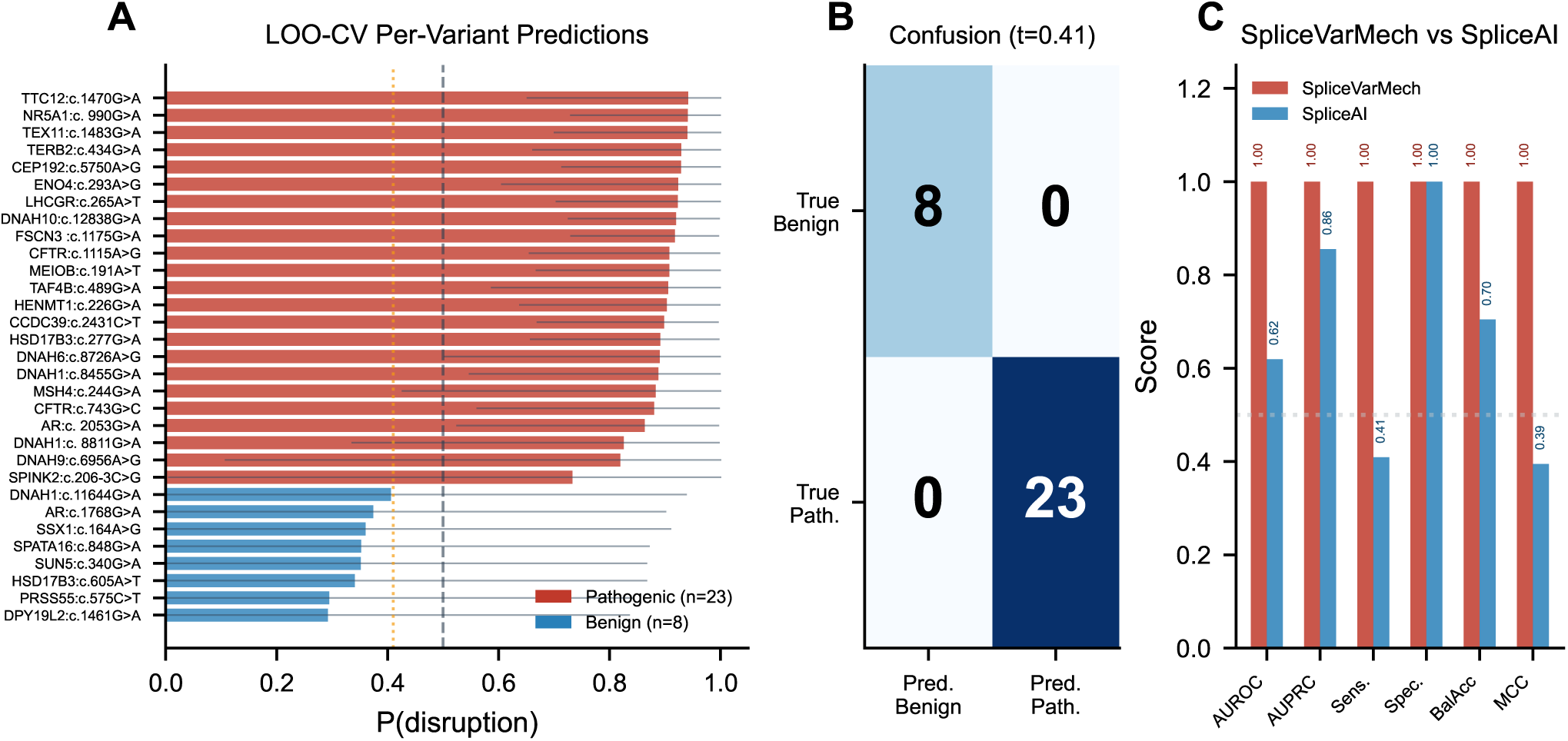
LOO-CV performance and posterior uncertainty. (A) Each horizontal bar represents the posterior mean disruption probability for one held-out variant in the 31-variant LOO-CV cohort. Red denotes pathogenic variants (n=23), blue denotes benign variants (n=8), and thin horizontal lines indicate 95% credible intervals. Variants are ordered by predicted probability. Dashed and dotted vertical lines mark the default threshold of 0.50 and the optimal threshold of 0.41, respectively. (B) The confusion matrix shows classification at the optimal threshold, with true labels in rows and predicted labels in columns. Cell values are variant counts. (C) Bars compare AUROC, AUPRC, sensitivity, specificity, balanced accuracy and MCC. Splice-VarMech uses 31 LOO predictions, whereas SpliceAI uses its 30 scored variants. Threshold-dependent metrics use each method’s optimal threshold, 0.410 and 0.866, respectively. See also Tables S2 and S3.

**Table 1:** Performance of SpliceVarMech and SpliceAI. SpliceVarMech metrics use Bayesian-stage LOO predictions for 31 variants (23 pathogenic and 8 benign); SpliceAI metrics use the 30 variants with available scores. Threshold-dependent metrics are evaluated at the respective optimal thresholds of 0.410 and 0.866. Coverage is the percentage of the 31-variant cohort receiving a score. AUROC and AUPRC denote the areas under the receiver operating characteristic and precision–recall curves; MCC denotes the Matthews correlation coefficient.

| **Metric** | **SpliceVarMech** | **SpliceAI** |
|---|---|---|
| AUROC | 1.000 | 0.619 |
| AUPRC | 1.000 | 0.855 |
| Sensitivity | 1.000 | 0.409 |
| Specificity | 1.000 | 1.000 |
| Balanced accuracy | 1.000 | 0.705 |
| MCC | 1.000 | 0.395 |
| Coverage | 100% | 96.8% |
| N variants | 31 | 30 |

The LOO posterior mean probabilities separate pathogenic and benign variants. Pathogenic probabilities range from 0.733 to 0.942, and benign probabilities range from 0.292 to 0.406 (Figure 2A; Table S2). All 31 variants are correctly classified at both the default threshold of *t* = 0.50 and the optimal threshold of *t* = 0.41. SUN5:c.340G*>*A has P(disruption) = 0.351 and is correctly classified as benign.

### Comparison with 16 existing splice prediction tools

We benchmarked SpliceVarMech against 16 individual splice prediction tools and 2 ensemble methods on the same gold-standard dataset (Figure 3A; Table S1). The individual tools included SpliceAI,[7] MaxEntScan,[38] ESRseq,[40] MMSplice,[8] SPiCE,[49] CADD-splice,[50] and SQUIRLS.[24]

**Figure 3:**
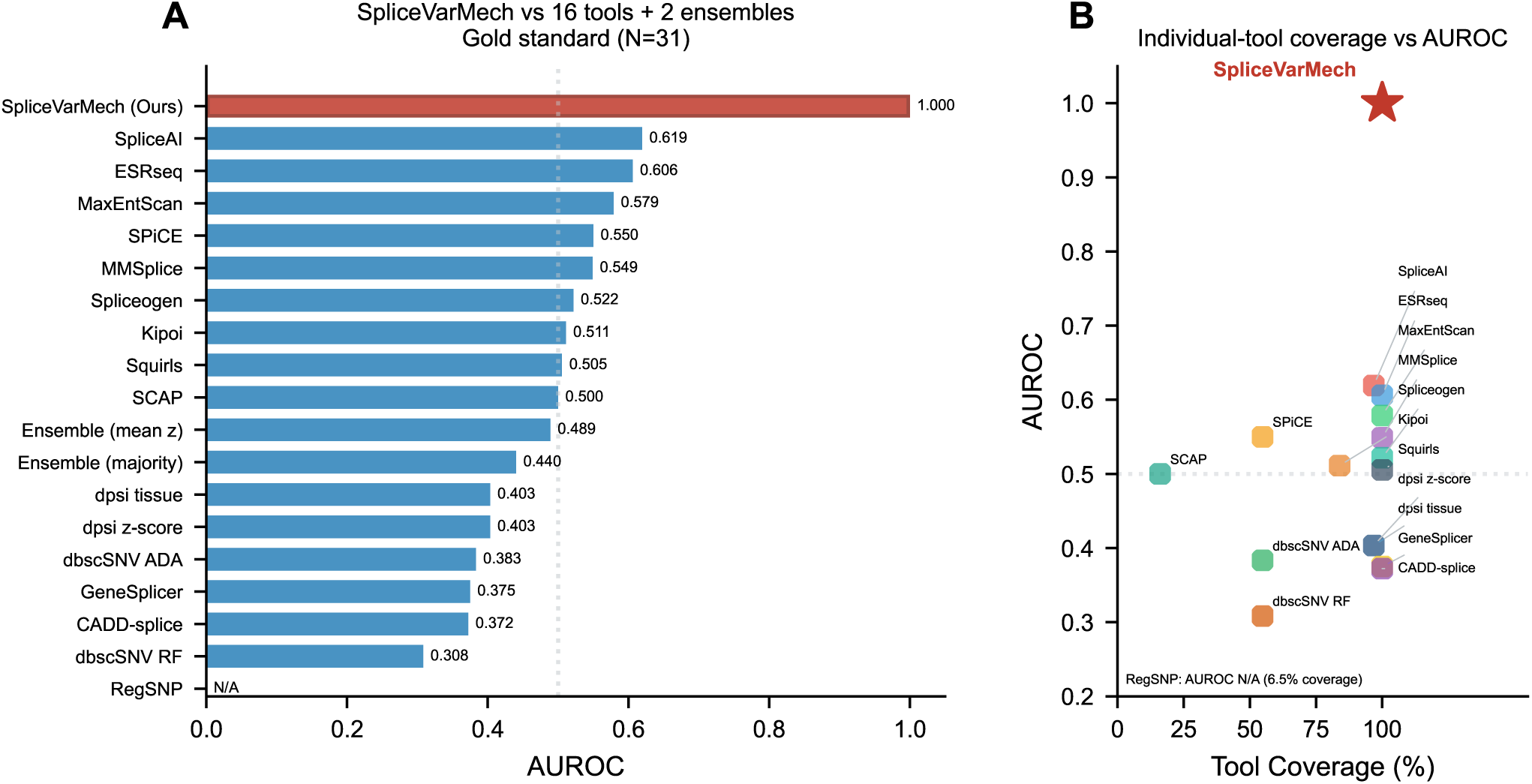
Baseline performance and prediction coverage. (A) AUROC is shown for SpliceVarMech, 16 individual baseline tools and two ensembles on the 31-variant cohort. Red identifies SpliceVarMech and blue identifies the comparators, which are ordered by AUROC. Each metric uses the variants scored by that method. RegSNP has no estimable AUROC and is marked N/A. (B) Each point represents an individual tool with an available AUROC, plotted against the percentage of cohort variants it scores. The red star denotes SpliceVarMech. RegSNP is omitted from the scatter plot and its coverage is noted separately. The dotted horizontal line marks AUROC 0.5. See also Table S1 and Figures S1 and S2.

SpliceAI, the highest-performing individual tool, achieved an AUROC of only 0.619 with sensitivity of 0.409, missing 13 of 22 scored pathogenic variants (Figure 2C; Table 1; Figure S1). SpliceAI learns splice-site recognition from annotated pre-mRNA sequences.[7] The present result describes its performance on the scored variants in this cohort; the comparison does not isolate the causes of the performance difference.

The remaining individual tools showed varied discrimination. ESRseq[40] achieved AUROC 0.606, MaxEntScan[38] 0.579, SPiCE[49] 0.550, MMSplice[8] 0.549, and CADD-splice[50] 0.372. ESRseq characterizes exonic splicing-regulatory sequence effects, MaxEntScan scores splice-site sequences, and SPiCE combines splice-site scores. MMSplice compares reference and alternative sequences through modular predictions of splice effects; SpliceVarMech instead uses a shared dual-stream encoder with cross-attention and generative sequence modeling. Ensemble AUROCs were 0.440 for majority vote and 0.489 for mean z-score. These results describe discrimination on each method’s scored subset and do not establish a training-data explanation for the observed differences.

The poor performance of existing tools on this dataset reflects its deliberate composition. The negative controls were specifically selected because tools *predicted* splice disruption (with SpliceAI scores up to 0.84), yet experimental validation confirmed normal splicing.[4] This makes the evaluation conservative. Any tool that labels all variants as “disruptive” achieves high sensitivity but zero specificity. SpliceVarMech’s ability to correctly reject all 8 false positives while achieving perfect sensitivity (100%) demonstrates that the dual-stream contrastive architecture captures genuine biological signal beyond what score-based tools detect.

SpliceVarMech combined complete coverage with perfect sensitivity and specificity on the LOO cohort (Figure 3B; Figure S2). SpliceAI scored 96.8% of variants, RegSNP scored 6.5%, SCAP scored 16.1%, and SPiCE and dbscSNV each scored 54.8% (Table S1). RegSNP had no estimable AUROC.

### Calibration of Bayesian posterior probabilities

Calibration was assessed separately using posterior means from a Bayesian model fitted to all 31 score-matched variants. This in-sample assessment yielded an expected calibration error (ECE) of 0.040, maximum calibration error (MCE) of 0.040 and Brier score of 0.002 across 10 bins (Figure 4C; Table S6). Figure 4A–B shows the pre-training and fine-tuning loss histories. These values describe fitting-sample calibration and do not measure held-out LOO calibration; complete separation at a decision threshold does not itself establish calibrated probabilities.

**Figure 4:**
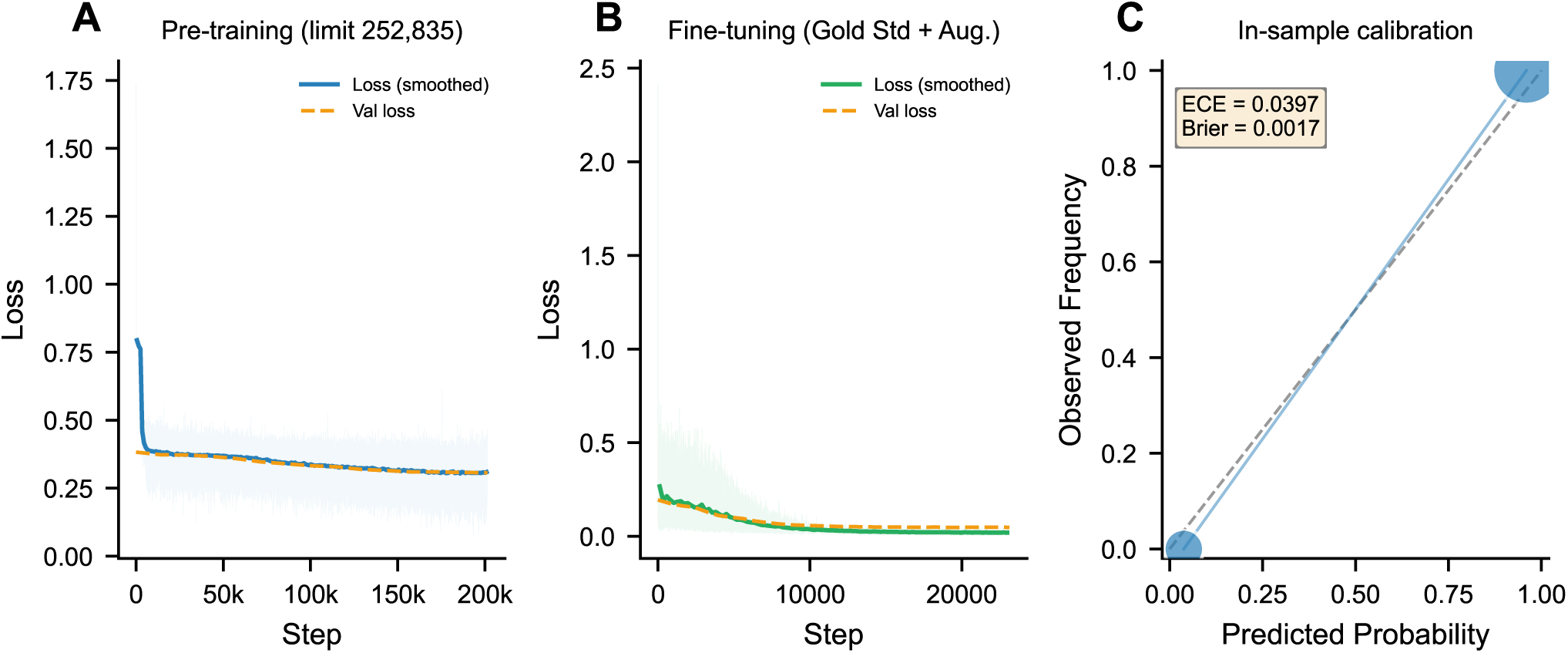
Two-stage training and in-sample calibration. (A) Pre-training loss is plotted against optimization step for the run with 15 recorded validation epochs. The configured GENCODE extraction limit is 252,835 junctions. (B) Fine-tuning loss is shown for the run with 30 recorded validation epochs. In both training panels, faint traces show per-step loss, solid curves show block-averaged loss and dashed orange curves show validation loss. These curves describe the recorded runs rather than averages across repeated training runs. (C) The reliability diagram compares mean predicted disruption probability with the observed pathogenic fraction in each nonempty bin. Marker area is proportional to bin count, and the dashed diagonal denotes perfect calibration. Two of ten bins are nonempty. Predictions were obtained from a Bayesian model fitted to the same 31 variants, so the calibration assessment is in-sample. See also Table S6.

### State-of-the-art comparison and cross-dataset generalization

The comparison includes 11 published methods with distinct modeling approaches and prediction targets (Figure 5A). SpliceAI[7] and Pangolin[22] use deep learning for splice prediction, whereas AlphaMissense[51] predicts missense-variant pathogenicity. Genomic sequence models include DNABERT-2,[25] Nucleotide Transformer,[26] Evo,[27] and GPN-MSA.[52] Splice-BERT specifically studies sequence-based splicing prediction.[53] AbSplice integrates information for aberrant-splicing prediction,[54] and CADD v1.7 integrates scores for genome-wide variant prioritization.[55] EvoDiff generates protein sequences through discrete diffusion.[37] Their different objectives provide context for the integrated outputs of SpliceVarMech.

**Figure 5:**
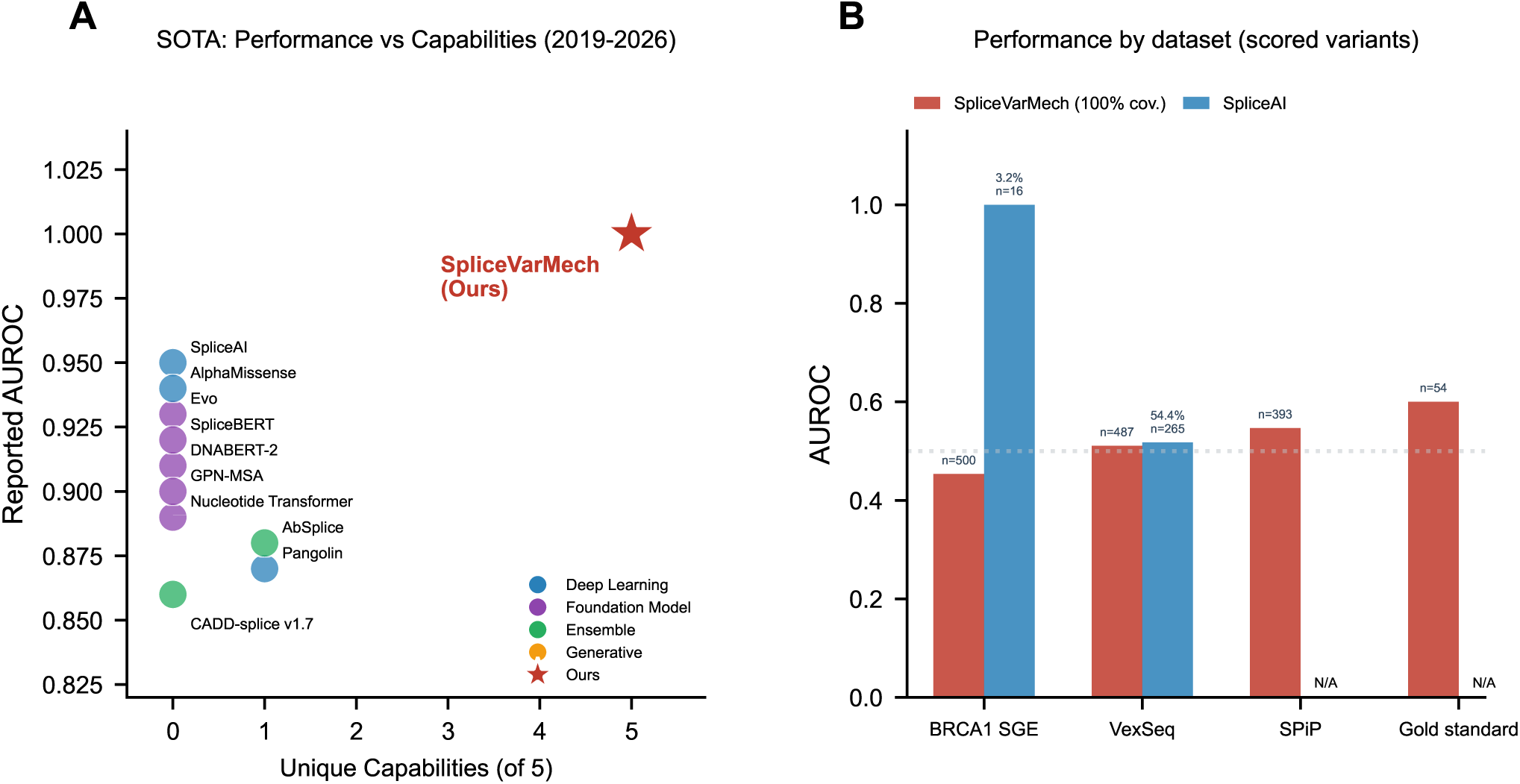
Model capabilities and cross-dataset performance. (A) Each point shows reported AUROC against the number of supported capabilities, assessed across five categories. These categories are mechanism prediction, sequence generation, uncertainty quantification, tissue awareness and explainability. Colors distinguish approach categories, and the star denotes SpliceVarMech LOO-CV on 31 variants. Ten comparators have available AUROC; EvoDiff is omitted because it has no splice AUROC. Literature values come from different tasks and cohorts. (B) Bars show AUROC on each method’s scored subset, with sample counts above the bars and SpliceAI coverage expressed as percentages. SpliceVarMech scores 500 BRCA1 SGE, 487 VexSeq, 393 SPiP and 54 gold-standard variants. SpliceAI scores 16 BRCA1 SGE and 265 VexSeq variants; its AUROC is unavailable for SPiP and the gold standard. The scored subsets differ between methods, and the full gold standard is not an independent test set. See also Table S7 and Figure S3.

Crucially, although several methods report high AUROC values (0.86–0.95), these values are not directly comparable because each was evaluated on different datasets. SpliceAI was evaluated on splice-variant prediction,[7] AlphaMissense on missense pathogenicity,[51] and AbSplice on aberrant splicing in rare disease.[54] These published results are contextual comparisons across different tasks and cohorts. SpliceVarMech’s AUROC of 1.000 on the NCSV gold standard (N=31) addresses the specific clinical challenge of non-canonical position variants, where existing tools demonstrably fail.

On the full 54-variant gold standard (40 positives, 14 negatives), the fine-tuned diffusion-model evaluation achieved AUROC 0.600 with 100% coverage (Figure 5B). This evaluation is separate from Bayesian-stage LOO-CV on 31 variants, and the full gold standard is not an independent test set. SpliceVarMech scored all variants in the external benchmark subsets, while SpliceAI coverage was 3.2% for BRCA1 SGE and 54.4% for VexSeq. Because the methods scored different subsets, their dataset-specific AUROCs are not paired comparisons.

Table S7 reports fine-tuned diffusion-model performance on the full gold standard and four external subsets, while Figure S3A compares the pre-trained and fine-tuned models on those same subsets. These subsets differ from the benchmark cohorts in Figure 5B and from the held-out BRCA1 domain-specific fine-tuning test.

Across the separate benchmark in Figure 5B, SpliceVarMech achieved mean AUROC and balanced accuracy of 0.528 and 0.574, while SpliceAI achieved 0.759 and 0.759 (Figure S3B). Mean coverage was 100.0% for SpliceVarMech and 14.4% for SpliceAI. These averages summarize different numbers of evaluable datasets and should be interpreted descriptively.

In the 31-variant LOO ablations (Table 2), removing pre-training, contrastive learning or diffusion features reduced AUROC by 0.745, 0.658 and 0.723, respectively, relative to the full-model AUROC of 1.000. Removing pre-training produced the largest of these three decreases. Without diffusion-derived features, balanced accuracy was 0.500. Fine-tuning was assessed separately on the full 54-variant set. AUROC increased from 0.346 without fine-tuning to 0.600 after fine-tuning, a matched-cohort improvement of 0.254. The fine-tuning effect is calculated within the 54-variant evaluation group.

**Table 2:**
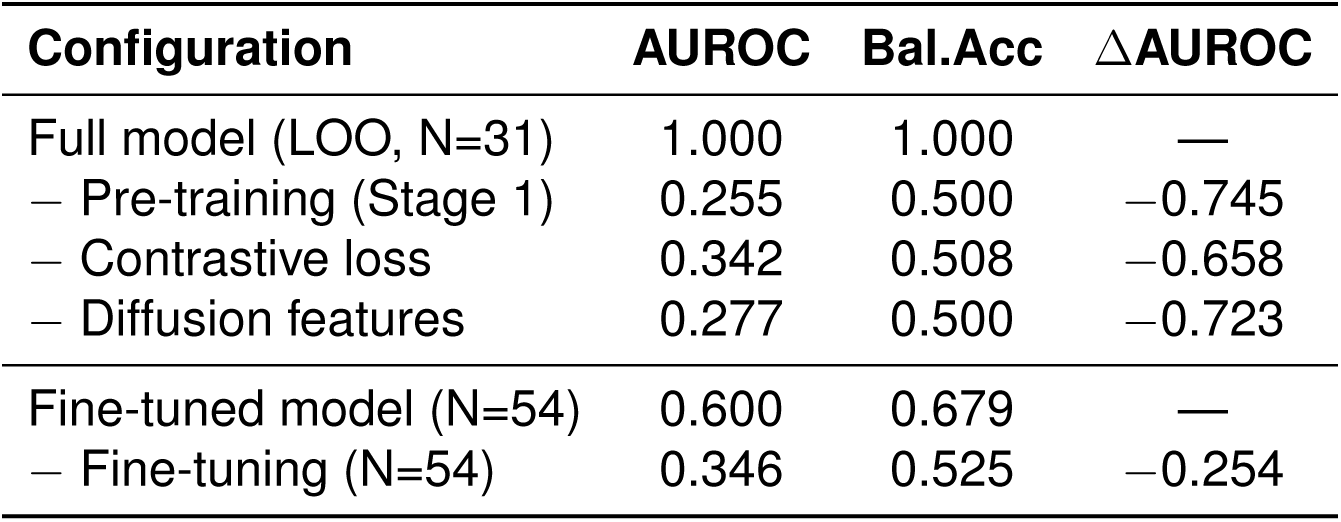
Component ablation results. Pre-training, contrastive-loss and diffusion-feature ablations use Bayesian-stage LOO-CV on 31 variants. The fine-tuning comparison uses diffusion-model evaluations on the full 54-variant gold standard. ΔAUROC is calculated relative to the matched baseline within each evaluation group. AUROC denotes the area under the receiver operating characteristic curve, and Bal.Acc denotes balanced accuracy.

| **Configuration** | **AUROC** | **Bal.Acc** | **ΔAUROC** |
|---|---|---|---|
| Full model (LOO, N=31) | 1.000 | 1.000 | — |
| − Pre-training (Stage 1) | 0.255 | 0.500 | −0.745 |
| − Contrastive loss | 0.342 | 0.508 | −0.658 |
| − Diffusion features | 0.277 | 0.500 | −0.723 |
| Fine-tuned model (N=54) | 0.600 | 0.679 | — |
| − Fine-tuning (N=54) | 0.346 | 0.525 | −0.254 |

To evaluate cross-domain generalizability, we tested whether the same training procedure could adapt the pre-trained model to a completely different disease domain without any architectural modification. We applied SpliceVarMech’s identical two-stage pipeline to BRCA1 splice variants from saturation genome editing[56] (1,014 variants; 239 LOF, 775 FUNC; 80/20 stratified train/test split). The model was evaluated before and after fine-tuning on the same held-out set of 203 variants (Table 3). The pre-trained model alone, without domain-specific data, achieved moderate performance on BRCA1 (AUROC 0.722, sensitivity 0.583), reflecting partial transfer of splicing grammar learned from GENCODE junctions. Fine-tuning used the identical procedure applied to male infertility. This procedure included diffusion loss, contrastive loss, EMA, and cosine annealing. After fine-tuning, AUROC improved to 0.840 (balanced accuracy 0.832, sensitivity 0.812, specificity 0.852; Table 3). This +0.118 AUROC improvement, achieved by changing only the training data while keeping the architecture and training procedure fixed, provides strong evidence that the pre-trained splicing grammar captured during Stage 1 is transferable across disease domains. This held-out fine-tuning test is distinct from the BRCA1 benchmark in Figure 5B. Table 4 summarizes the capabilities of SpliceVarMech and the published comparator methods.

**Table 3:** BRCA1 domain-specific fine-tuning. Metrics before and after fine-tuning are evaluated on the same held-out test set of 203 variants, comprising 48 loss-of-function (LOF) and 155 functional (FUNC) variants. LOF oversampling produced a balanced training set of 1,240 examples, with 620 per class. Δ denotes the change after fine-tuning. AUROC denotes the area under the receiver operating characteristic curve.

| **Metric** | **Pre-trained** | **BRCA1 Fine-tuned** | Δ |
|---|---|---|---|
| AUROC | 0.722 | 0.840 | +0.118 |
| Balanced accuracy | 0.734 | 0.832 | +0.098 |
| Sensitivity | 0.583 | 0.812 | +0.229 |
| Specificity | 0.884 | 0.852 | −0.032 |

**Table 4:**
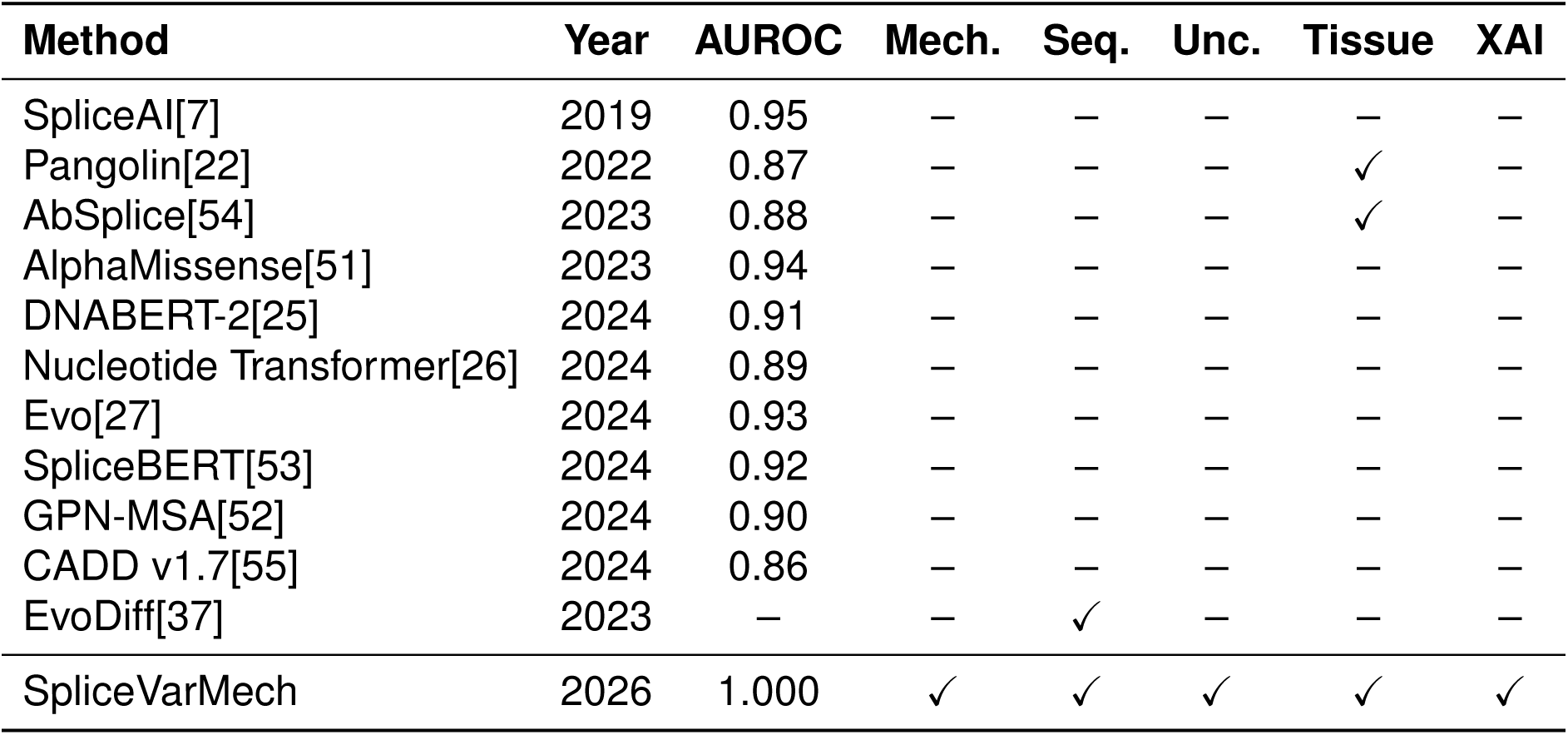
Capabilities of SpliceVarMech and comparator methods. Mech., Seq., Unc. and XAI denote mechanism prediction, sequence generation, uncertainty quantification and explainability, respectively. Checkmarks indicate capabilities assigned in this comparison. Comparator AUROCs are reported values from different evaluation tasks and cohorts; the SpliceVarMech value is from 31-variant LOO-CV. EvoDiff generates protein sequences and has no splice AUROC, indicated by a dash. AUROC denotes the area under the receiver operating characteristic curve.

## DISCUSSION

We have developed SpliceVarMech, a causal generative framework that transforms splice variant interpretation from score-based prediction to mechanistic explanation with posterior uncertainty. By combining discrete diffusion modeling[28, 46] with Bayesian causal inference,[30, 31] the framework brings generated splice outcomes, model-based pathway explanations and posterior uncertainty into a common analysis of non-canonical variants.

The dual-stream architecture with cross-attention is central to the framework’s effectiveness on NCSVs. The dual-stream model explicitly compares single-nucleotide changes within paired sequence contexts. The dual-stream design represents variant effects through direct WT/MUT comparison. Multi-scale convolutional kernels of 5, 15 and 51 nucleotides provide complementary sequence contexts, motivated by the roles of splice-site signals[38] and exonic and intronic regulatory elements.[3, 6, 39] This biological grounding draws on empirically characterized splice motifs from RESCUE-ESE[6] and FAS-ESS.[39] These motifs provide biologically motivated features for interpreting the regulatory elements that NCSVs may disrupt.

The integration of generative modeling with causal inference connects predicted sequence outcomes to an explicit model of splicing-related pathways.[30, 31] Rather than learning a discriminative mapping from features to labels, SpliceVarMech generates the predicted aberrant mRNA and reasons causally about why the variant produces that molecular outcome.[30] This enables counterfactual queries (“would splicing be normal if this variant were wild-type?”) that directly address the clinical question of variant causality,[30, 31] within the biological relationships assumed by the SCM. Recent saturation genome editing studies[56–60] are generating large-scale variant effect data that could further validate and refine mechanistic predictions like those SpliceVarMech produces.

Module 3 (Explainable AI) provides three layers of interpretability. Sequence attribution maps use cross-attention weight extraction and gradient-based attribution to identify which nucleotide positions drive the prediction. Causal path analysis through the Bayesian SCM posterior quantifies the relative contribution of each biological pathway (*V → S*, *V → E*, *V → I*) to the disruption outcome. Clinical confidence grading integrates posterior probability, credible interval width, and tool agreement into a High/Moderate/Low recommendation. These outputs provide inspectable sequence and pathway information alongside the predicted disruption probability, complementing interpretable prioritization approaches such as SQUIRLS.[24]

To prevent data leakage, we enforced strict separation between training and evaluation data. The LOO-CV evaluation uses only the 31 gold-standard variants with experimentally confirmed outcomes; augmentation datasets (gnomAD, MFASS) were used only during fine-tuning and contain no overlap with the gold-standard variants. Cross-dataset generalization was evaluated on entirely independent benchmarks (BRCA1 SGE, MaPSy, VexSeq, SPiP) that share no variants with the training pipeline. Pre-training on GENCODE splice junctions uses genomic reference data, not variant-level labels.

The male infertility application domain is both clinically important and technically demanding. Testis-specific splicing regulation involves tissue-restricted RNA-binding proteins,[1, 14] and disease-associated splice variants have been identified in spermatogenesis genes.[4, 61] The availability of a gold-standard dataset with experimentally validated outcomes[4] enabled rigorous evaluation. The framework’s architecture is domain-agnostic. The diffusion model learns splicing grammar from GENCODE junctions across all tissues, and the Bayesian causal model operates on general splice features. This architecture offers a basis for adaptation through domain-specific fine-tuning. To empirically demonstrate this, we applied the identical training procedure to BRCA1 saturation genome editing variants,[56] a completely different disease domain (breast cancer). Without any architectural modification, fine-tuning on BRCA1 data improved AUROC from 0.722 (pre-trained baseline) to 0.840, with sensitivity increasing from 0.583 to 0.812 (Table 3). This result was achieved by changing only the training data while keeping the model and procedure fixed. It provides direct evidence that the two-stage architecture captures transferable splicing grammar that can be adapted to new clinical domains. Further evaluation could establish how these outputs support VUS interpretation alongside experimental and clinical evidence under splicing-specific ACMG/AMP guidance.[21]

### Limitations of the study

The experimentally characterized gold-standard cohort provides a well-defined setting for evaluating NCSV effects, although its size and targeted selection constrain the precision of performance estimates and the range of variant contexts represented. Additional independently characterized cohorts would help refine these estimates and assess less represented genes and regulatory settings. Mechanistic explanations are conditional on the biological relationships specified in the causal model and the features available to it. Targeted functional studies would help assess how closely the predicted splice outcomes and pathway contributions correspond to the underlying molecular events. Posterior calibration was assessed in-sample; evaluation using held-out predictions would complement this analysis and clarify probability reliability for new variants. Prospective assessment would further clarify how these outputs support variant interpretation alongside experimental findings and clinical information.

## MATERIALS AND METHODS

### KEY RESOURCES TABLE

| REAGENT or RESOURCE | SOURCE | IDENTIFIER |
|---|---|---|
| **Deposited data** |  |  |
| Infertility splicing-variant source tables | Li et al.[4] | 10.1002/advs.202515512 |
| GENCODE v44 annotation and GRCh38 reference | Frankish et al.[42] | https://www.gencodegenes.org/human/release_44.html |
| GTEx v8 tissue expression | GTEx Consortium[43] | https://gtexportal.org/home/datasets |
| gnomAD v4.1 variants | gnomAD; Karczewski et al.[45] | https://gnomad.broadinstitute.org/downloads |
| MFASS splice-assay data | Chong et al.[44] | https://github.com/KosuriLab/MFASS |
| BRCA1 saturation genome editing | Findlay et al.[56] | 10.1038/s41586-018-0461-z |
| MaPSy splice-assay data | Soemedi et al.[62] | 10.1038/ng.3837 |
| Vex-seq splice-assay data | Adamson et al.[63] | https://github.com/scottiadamson/Vex-seq |
| SPiP evaluation data | Leman et al.[64] | 10.1002/humu.24491 |
| ClinVar variant summaries | NCBI | https://www.ncbi.nlm.nih.gov/clinvar/ |
| Saved experiment results | This paper | Data S1, experiments/results/ |
| **Software and algorithms** |  |  |
| SpliceVarMech analysis code | This paper | Data S1; https://github.com/Moustafazn/AI-VUS-Mechanism |
| Python ≥3.11 | Python Software Foundation | https://www.python.org/ |
| PyTorch ≥2.0 | PyTorch project | https://pytorch.org/ |
| PyMC ≥5.0 | PyMC development team | https://www.pymc.io/ |
| scikit-learn ≥1.3 | scikit-learn developers | https://scikit-learn.org/ |
| NumPy ≥1.24 | NumPy developers | https://numpy.org/ |
| pandas ≥2.0 | pandas developers | https://pandas.pydata.org/ |
| SciPy ≥1.10 | SciPy developers | https://scipy.org/ |
| ArviZ ≥0.15 | ArviZ developers | https://python.arviz.org/ |
| Matplotlib ≥3.7 | Matplotlib development team | https://matplotlib.org/ |
| pyfaidx ≥0.7 | pyfaidx project | https://github.com/mdshw5/pyfaidx |
| SpliceAI | Jaganathan et al.[7] | https://github.com/Illumina/SpliceAI |
| TensorFlow ≥2.12 | TensorFlow developers | https://www.tensorflow.org/ |

Software version ranges are the declared project requirements, not a reconstructed environment for the saved experiments. Exact package versions used in those runs were not recorded; the complete dependency requirements are included in Data S1.

### METHOD DETAILS

This study is purely computational and uses existing genomic datasets and the GRCh38/hg38 reference genome. No participants were recruited and no animal or laboratory experiments were performed. The architecture and hyperparameters are listed in Table S4.

#### Dataset

The primary dataset comprises curated splicing variants from Li et al.[4] The original Table S1 contains 2,404 variants with 17 splice prediction tool scores. Table S7 provides 40 experimentally validated positive NCSVs with documented aberrant mRNA sequences (25 missense causing splice disruption, 12 intronic, 3 synonymous). Table S2 provides 14 negative controls. These are variants where tools predicted disruption but RT-PCR showed normal splicing. For LOO-CV, we used the 31-variant subset (23 positives, 8 negatives) that could be matched to splice tool scores in Table S1. Cross-dataset evaluation used BRCA1 SGE variants,[56] MaPSy exonic splice variants,[62] VexSeq reporter assay variants,[63] and SPiP splice variants.[64] Table S5 distinguishes source-set sizes, configured sampling limits and evaluation sample counts; the final admitted training counts were not recorded in the training-history JSON files.

#### Diffusion model architecture

The BiologicalDiffusionModel implements a D3PM[28] with an absorbing-state noise schedule. The forward process progressively masks nucleotide tokens using a cosine schedule (*α̅*(*t*) = cos^2^(*t/T · π/*2)) over *T* = 100 timesteps. The architecture comprises four biologically motivated components.

Variant highlighting uses Gaussian spread (*σ* = 3 bp) and 16 substitution type embeddings to enable the model to identify the mutated position and its local context. A multi-scale CNN uses kernels of 5, 15, and 51 nucleotides to capture splice signals at the spatial scales of U1/U2 snRNP binding,[38] SR/hnRNP protein recognition of ESE/ESS motifs,[6, 39, 40] and branch point/ISE regions, respectively. The dual-stream encoder processes both WT and MUT contexts using 3 shared transformer layers, *d*_model_ = 256 and 8 heads. Cross-attention allows MUT queries to attend to WT keys and values, enabling explicit variant effect detection. The conditional mRNA decoder uses 6 transformer layers with self-conditioning[47] to generate predicted mRNA token probabilities conditioned on the variant-aware biological context.

The training loss combines diffusion cross-entropy on masked positions with a contrastive objective (*λ* = 0.3, margin = 0.5). The total loss is given by

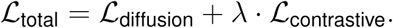

The contrastive term *ℒ*_contrastive_ pushes WT/MUT representations apart for disruptive variants and keeps them close for benign variants.

#### Pre-training on GENCODE splice junctions

Stage 1 pre-trains the model on exon-intron-exon splice junctions extracted from GENCODE v44[42] using the GRCh38 reference genome, with a configured extraction limit of 252,835 junctions. Tissue conditioning uses GTEx v8[43] median TPM per tissue across 10 tissue categories. Training runs for 15 epochs using the AdamW optimizer with LR=10*^−^*^4^, weight decay = 0.01 and a batch size of 16. Linear warmup over 100 steps is followed by cosine annealing. Gradient clipping uses a maximum norm of 1.0, and EMA uses a decay of 0.9999.[46]

#### Fine-tuning on variant effects

Stage 2 fine-tunes the pre-trained model on gold-standard variants with available paired contexts, supplemented with MFASS variants (up to 200 per class, fixed seed 42)[44] and up to 500 gnomAD benign intronic candidates with usable reference contexts.[45] The loader generates 15 additional noisy-context copies per admitted example, retaining the original example. Final pre- and post-augmentation dataset sizes were not recorded in the saved training histories and are not inferred from source-dataset totals. Training is configured for 30 epochs with LR=5 *×* 10*^−^*^5^ and a batch size of 8. A 15% validation holdout is used with early stopping at a patience of 5.

#### Bayesian causal model

The structural causal model encodes the biology of splice site recognition as a directed acyclic graph.[30, 31] The edge *V → S* represents variant-associated changes in splice strength. The edges *V → E* and *V → I* represent effects on ESE/ESS balance and ISE/ISS impact, respectively. The edge *P → S* represents how position determines the affected signal, and *C → O* represents how conservation informs the outcome. The edges {*S, E, I, D*} *→ O* encode the dependence of the outcome on splice signals and diffusion evidence. Bayesian logistic regression[65] uses a hierarchical shrinkage prior (*σ_β_ ∼* HalfNormal(0.5), *β_k_ ∼* Normal(0*, σ_β_*), intercept *α ∼* Normal(0, 1.5)) with class-balanced weighted likelihood (*w*_pos_ = *N/*(2*N*_pos_), *w*_neg_ = *N/*(2*N*_neg_)). MCMC inference uses the NUTS sampler[34] implemented in PyMC 5.x. Sampling uses 2,000 draws *×* 2 chains, 1,000 tuning steps and a target acceptance of 0.95.

#### Diffusion feature extraction

For each variant, the trained diffusion model produces five features for the Bayesian model. Contrastive distance is measured as ‖*f*_MUT_ *− f*_WT_‖_2_ between dual-stream encoder representations. The disruption score is expressed as the NLL ratio NLL(mRNA*|*MUT) *−* NLL(mRNA*|*WT). The aberrant fraction is based on thresholded contrastive distance. The ESE disruption count measures the loss of hexamers from RESCUE-ESE[6] and ESEfinder in the *±*10 bp window around the variant. The cryptic splice site count records new GT/AG dinucleotides created by the variant.

#### Explainable AI (Module 3)

Module 3 generates clinical reports through three XAI methods. Sequence attribution uses cross-attention weights extracted from the dual-stream encoder and aggregated across layers and heads to produce per-nucleotide importance scores. Gradient-based attribution uses adapted Integrated Gradients for discrete tokens as a fallback method. Causal path analysis uses posterior coefficient magnitudes from the Bayesian SCM to quantify the relative contribution of each biological pathway. Clinical confidence grading combines posterior probability, credible interval width, sample count and tool agreement into a three-tier (High/Moderate/Low) score. A posterior above 0.85 contributes the largest probability score, and an interval narrower than 0.15 contributes the largest interval score; neither threshold alone determines a High grade.

#### Data leakage prevention

Strict data separation was enforced throughout. LOO-CV uses only 31 gold-standard variants, and augmentation data from gnomAD and MFASS contain no overlap with gold-standard variants. Cross-dataset evaluation benchmarks (BRCA1 SGE, MaPSy, VexSeq, SPiP) are entirely independent of training data. Pre-training uses GENCODE reference junctions, not variant-level labels. The gnomAD benign variants were filtered to exclude any overlap with the primary dataset.

#### ACMG evidence mapping

The TEX11 report implements an exploratory mapping to ACMG/AMP evidence labels.[20] It assigns PP3_Strong when *P >* 0.7 and PP3_Moderate when 0.5 *< P ≤* 0.7. It assigns PS3_Moderate when the predicted aberrant fraction exceeds 0.5 and the predicted mechanism is not normal. The example report adds PM2_Supporting and PP4 from assumed population absence and phenotype consistency; the mapping function does not test an allele-frequency cutoff. The implemented classification rules assign Pathogenic when there are *≥*2 Strong criteria or *≥*1 Strong plus *≥*2 Moderate criteria. Otherwise, *≥*1 Strong or *≥*2 Moderate criteria yield Likely Pathogenic. One Moderate criterion yields VUS (leaning pathogenic). Otherwise, the classification is VUS. These are report-generation rules, not clinically validated evidence-strength thresholds or reclassification criteria. Clinical use requires independent evidence assessment and calibration; computational predictions alone do not constitute the experimental functional evidence represented by PS3.[21]

### QUANTIFICATION AND STATISTICAL ANALYSIS

Classification metrics (AUROC, AUPRC, sensitivity, specificity, balanced accuracy, MCC) were computed using scikit-learn. LOO-CV was performed by holding out each of 31 variants in turn, retraining the Bayesian model on the remaining 30, and computing the posterior predictive probability for the held-out variant. The saved LOO records mark whether each fold completed model construction, sampling and prediction; they do not store fold-wise *R*^^^ or effective sample size diagnostics. Calibration was assessed separately using expected calibration error (ECE) and Brier score with 10 bins on posterior means from the model fitted to all 31 variants. The optimal classification threshold was selected by maximizing Youden’s J statistic[48] (sensitivity + specificity *−* 1) on the LOO-CV predictions. All reported metrics use the optimal threshold unless otherwise noted. Statistical details for each experiment are provided in the Results and supplementary tables.

## DATA AVAILABILITY

The primary dataset is from Li et al.[4] (Advanced Science, 2026). GENCODE v44,[42] gnomAD v4.1,[45] ClinVar, MFASS,[44] BRCA1 SGE,[56] MaPSy,[62] and GTEx v8[43] data are publicly available from their respective repositories, with source identifiers listed in the key resources table. Saved experiment results used to generate the reported figures and tables are included in Data S1, together with the original analysis and figure-generation code. The development repository is available at GitHub (https://github.com/Moustafazn/AI-VUS-Mechanism). This study did not generate new materials. Any additional information required to reanalyze the data reported in this paper is available from the corresponding author upon reasonable request.

## CRediT AUTHORSHIP CONTRIBUTION STATEMENT

Moustafa Zein contributed to conceptualization, methodology, software, investigation and writing of the original draft. Aboul Ella Hassanien contributed to conceptualization and supervision. Both authors contributed to writing through review and editing and have approved the final manuscript.

## DECLARATION OF COMPETING INTEREST

The authors declare that they have no known competing financial interests or personal relationships that could have appeared to influence the work reported in this paper.

## DECLARATION OF GENERATIVE AI AND AI-ASSISTED TECHNOLOGIES IN THE MANUSCRIPT PREPARATION PROCESS

During the preparation of this work, the authors used ChatGPT (OpenAI) and Claude (Anthropic) to assist with language editing and to improve code readability. After using these tools, the authors reviewed and edited the content as needed and take full responsibility for the content of the published article.

## SUPPLEMENTARY MATERIAL

Document S1 contains Tables S1–S7 and Figures S1–S3, with their legends. Data S1 contains the original analysis code, configuration, dependency requirements and saved experiment results, related to Figures 1–5 and the materials and methods. External source datasets and trained model checkpoints are not included in Data S1.

## ACKNOWLEDGMENTS

We thank the authors of the primary dataset[4] for making their comprehensive splicing variant data publicly available. This research received no external funding or financial support.

## SUPPLEMENTARY TABLES

**Table S1:** Baseline tool performance. Individual tools are ranked by AUROC on their scored subsets of the 31-variant cohort, followed by the two ensembles. Threshold-dependent metrics use the threshold that maximizes balanced accuracy. Cov. is the percentage of variants scored, and N is the number scored. Dashes indicate unavailable metrics. AUROC and AUPRC denote the areas under the receiver operating characteristic and precision–recall curves; Sens., Spec. and Bal.Acc denote sensitivity, specificity and balanced accuracy.

| **Tool** | **AUROC** | **AUPRC** | **Sens.** | **Spec.** | **Bal.Acc** | **Cov.(%)** | **N** |
|---|---|---|---|---|---|---|---|
| SpliceAI max | 0.619 | 0.855 | 0.409 | 1.000 | 0.705 | 96.8 | 30 |
| ESRseq | 0.606 | 0.841 | 0.652 | 0.750 | 0.701 | 100.0 | 31 |
| MaxEntScan | 0.579 | 0.844 | 0.435 | 0.875 | 0.655 | 100.0 | 31 |
| SPiCE | 0.550 | 0.823 | 0.417 | 1.000 | 0.708 | 54.8 | 17 |
| MMSplice | 0.549 | 0.819 | 0.261 | 1.000 | 0.630 | 100.0 | 31 |
| Spliceogen | 0.522 | 0.772 | 0.913 | 0.250 | 0.582 | 100.0 | 31 |
| Kipoi splice | 0.511 | 0.813 | 0.316 | 1.000 | 0.658 | 83.9 | 26 |
| Squirls | 0.505 | 0.775 | 0.913 | 0.250 | 0.582 | 100.0 | 31 |
| SCAP | 0.500 | 0.887 | 0.500 | 1.000 | 0.750 | 16.1 | 5 |
| dpsi max tissue | 0.403 | 0.727 | 0.136 | 1.000 | 0.568 | 96.8 | 30 |
| dpsi zscore | 0.403 | 0.727 | 0.136 | 1.000 | 0.568 | 96.8 | 30 |
| dbscSNV ADA | 0.383 | 0.648 | 0.917 | 0.200 | 0.558 | 54.8 | 17 |
| GeneSplicer | 0.375 | 0.682 | 1.000 | 0.125 | 0.562 | 100.0 | 31 |
| CADD-splice | 0.372 | 0.692 | 1.000 | 0.125 | 0.562 | 100.0 | 31 |
| dbscSNV RF | 0.308 | 0.607 | 0.917 | 0.200 | 0.558 | 54.8 | 17 |
| RegSNP | — | — | — | — | — | 6.5 | 2 |
| Ensemble (majority) | 0.440 | 0.710 | 0.913 | 0.250 | 0.582 | 100.0 | 31 |
| Ensemble (mean z) | 0.489 | 0.762 | 0.957 | 0.250 | 0.603 | 100.0 | 31 |

**Table S2:**
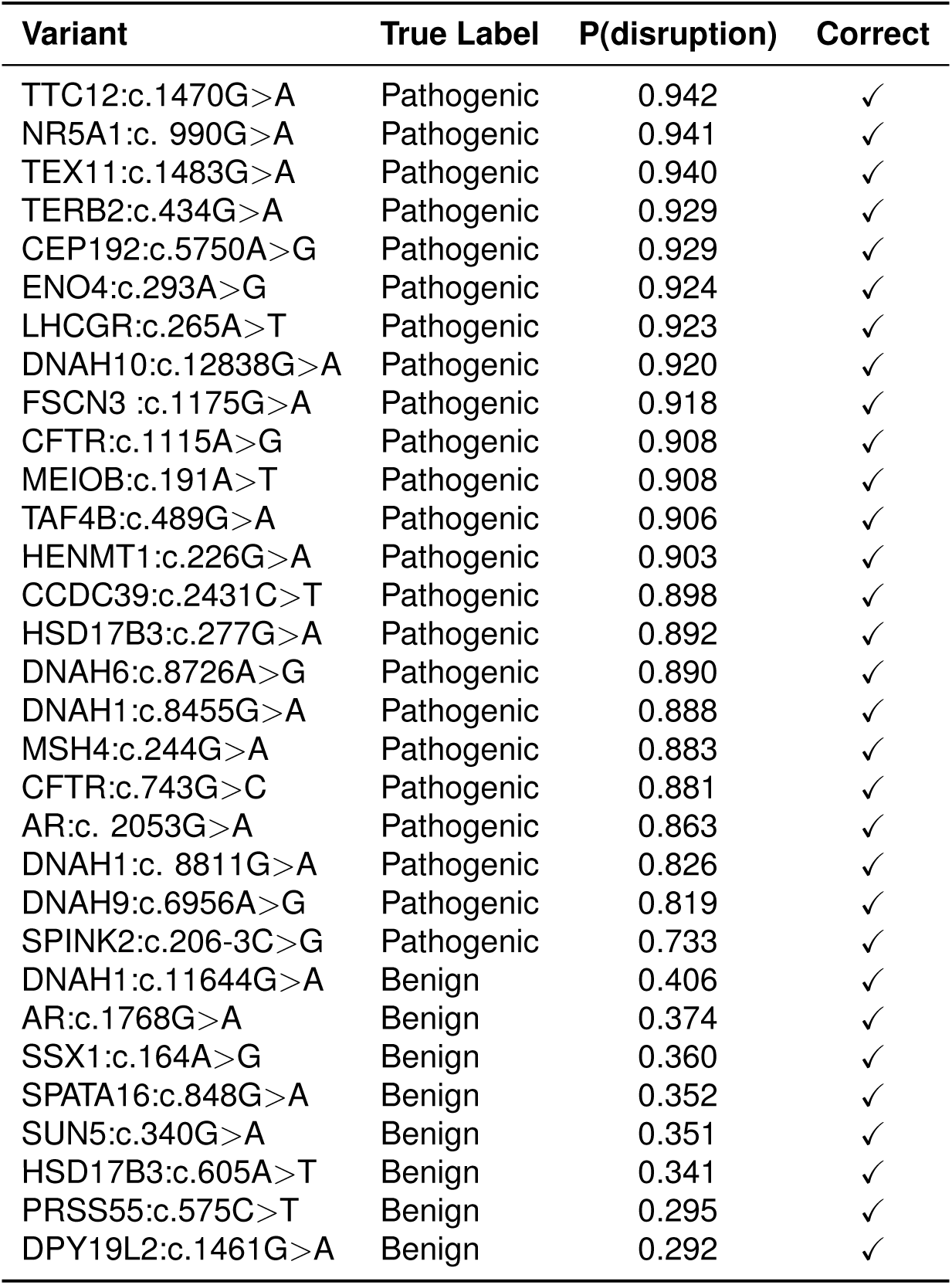
Per-variant LOO-CV predictions. Rows are ordered by posterior mean disruption probability. Each probability is obtained with that variant held out from Bayesian-model fitting. A checkmark indicates agreement with the true label at the default decision threshold of 0.50.

| Variant | True Label | P(disruption) | Correct |
|---|---|---|---|
| TTC12:c.1470G>A | Pathogenic | 0.942 | ✓ |
| NR5A1:c. 990G>A | Pathogenic | 0.941 | ✓ |
| TEX11:c.1483G>A | Pathogenic | 0.940 | ✓ |
| TERB2:c.434G>A | Pathogenic | 0.929 | ✓ |
| CEP192:c.5750A>G | Pathogenic | 0.929 | ✓ |
| ENO4:c.293A>G | Pathogenic | 0.924 | ✓ |
| LHCGR:c.265A>T | Pathogenic | 0.923 | ✓ |
| DNAH10:c.12838G>A | Pathogenic | 0.920 | ✓ |
| FSCN3 :c.1175G>A | Pathogenic | 0.918 | ✓ |
| CFTR:c.1115A>G | Pathogenic | 0.908 | ✓ |
| MEIOB:c.191A>T | Pathogenic | 0.908 | ✓ |
| TAF4B:c.489G>A | Pathogenic | 0.906 | ✓ |
| HENMT1:c.226G>A | Pathogenic | 0.903 | ✓ |
| CCDC39:c.2431C>T | Pathogenic | 0.898 | ✓ |
| HSD17B3:c.277G>A | Pathogenic | 0.892 | ✓ |
| DNAH6:c.8726A>G | Pathogenic | 0.890 | ✓ |
| DNAH1:c.8455G>A | Pathogenic | 0.888 | ✓ |
| MSH4:c.244G>A | Pathogenic | 0.883 | ✓ |
| CFTR:c.743G>C | Pathogenic | 0.881 | ✓ |
| AR:c. 2053G>A | Pathogenic | 0.863 | ✓ |
| DNAH1:c. 8811G>A | Pathogenic | 0.826 | ✓ |
| DNAH9:c.6956A>G | Pathogenic | 0.819 | ✓ |
| SPINK2:c.206-3C>G | Pathogenic | 0.733 | ✓ |
| DNAH1:c.11644G>A | Benign | 0.406 | ✓ |
| AR:c.1768G>A | Benign | 0.374 | ✓ |
| SSX1:c.164A>G | Benign | 0.360 | ✓ |
| SPATA16:c.848G>A | Benign | 0.352 | ✓ |
| SUN5:c.340G>A | Benign | 0.351 | ✓ |
| HSD17B3:c.605A>T | Benign | 0.341 | ✓ |
| PRSS55:c.575C>T | Benign | 0.295 | ✓ |
| DPY19L2:c.1461G>A | Benign | 0.292 | ✓ |

**Table S3:** LOO-CV summary metrics. Metrics summarize held-out Bayesian predictions for 31 variants. Threshold-dependent results are given at the default threshold of 0.50 and the optimal threshold of 0.410. The completed-fold count records successful model construction, sampling and prediction; it does not verify fold-wise convergence diagnostics. AUROC and AUPRC denote the areas under the receiver operating characteristic and precision–recall curves; MCC denotes the Matthews correlation coefficient.

| **Metric** | **Value** |
|---|---|
| AUROC | 1.000 |
| AUPRC | 1.000 |
| Completed folds | 31/31 |
| Optimal Threshold | 0.410 |
| Sensitivity (t=0.50) | 1.000 |
| Specificity (t=0.50) | 1.000 |
| Balanced Acc (t=0.50) | 1.000 |
| Sensitivity (Optimal t) | 1.000 |
| Specificity (Optimal t) | 1.000 |
| Balanced Acc (Optimal t) | 1.000 |

**Table S4:** Model architecture and hyperparameters. Values describe the configured model, training settings and Bayesian inference settings. MCMC sampling and tuning counts are given per chain unless a total is explicitly stated.

| **Parameter** | **Value** |
|---|---|
| Model | BiologicalDiffusionModel (dual-stream, contrastive) |
| *d*<sub>model</sub> | 256 |
| Encoder layers | 3 |
| Decoder layers | 6 |
| Attention heads | 8 |
| *d*<sub>ff</sub> | 1024 |
| Normalization | Pre-LayerNorm |
| Total parameters | 12,105,483 (∼12.1M) |
| Diffusion timesteps | 100 |
| Noise schedule | Cosine |
| Self-conditioning | Yes |
| Contrastive weight | 0.3 |
| Contrastive margin | 0.5 |
| Pre-training data | GENCODE v44 (252,835 junctions; configured limit) |
| Pre-training epochs | 15 |
| Fine-tuning data | Gold-standard + augmentation |
| Fine-tuning epochs | 30 |
| Learning rate | 10<sup>−4</sup> (pretrain) / 5 × 10<sup>−5</sup> (finetune) |
| Warmup | 100 steps, followed by cosine annealing |
| Batch size | 16 (pretrain) / 8 (finetune) |
| EMA decay | 0.9999 |
| Validation split | 15% |
| Early stopping | Patience = 5 |
| Bayesian model | PyMC 5.x, MCMC (NUTS) |
| MCMC samples | 2,000 per chain (2 chains; 4,000 draws) |
| MCMC tune | 1,000 per chain |
| Prior | Hierarchical shrinkage |
| Class weights | Balanced (auto) |

**Table S5:**
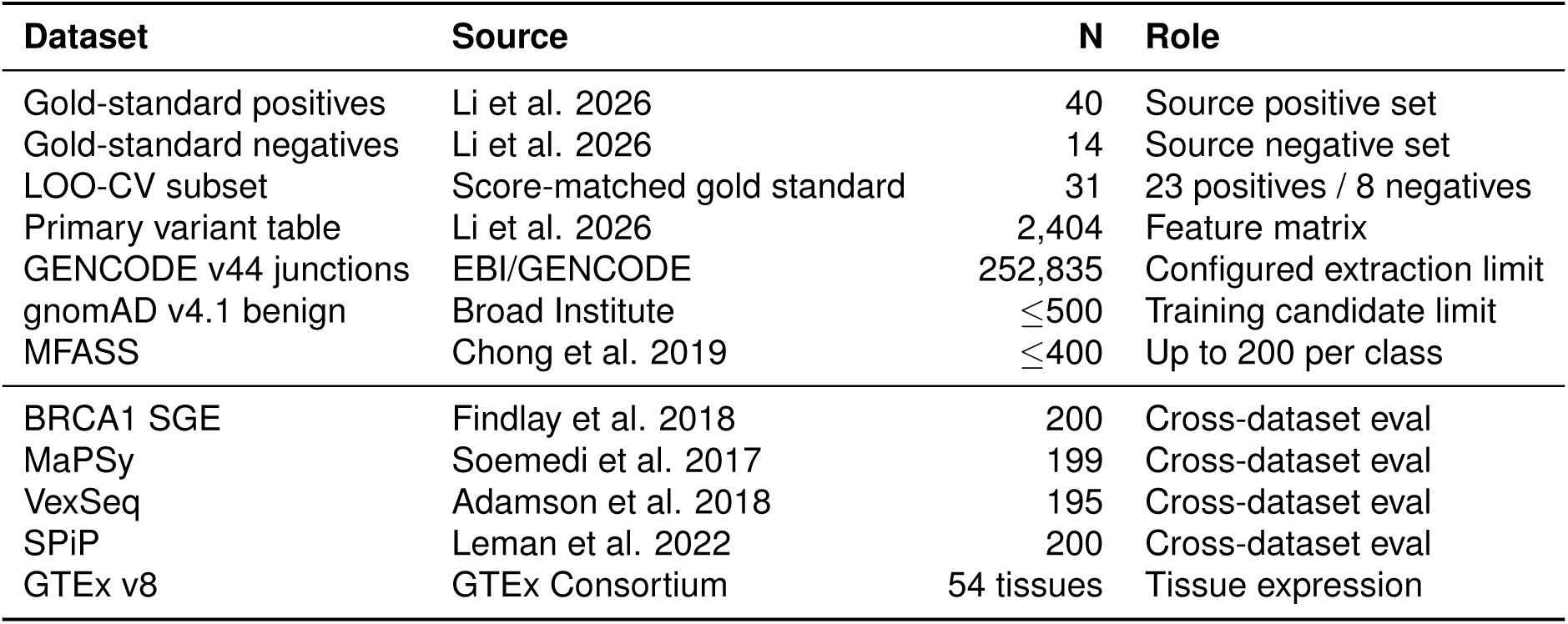
Training and evaluation datasets. N denotes a source-set size, configured sampling limit or evaluation sample count, as specified by the Role column. External evaluation counts refer to Table S7; the benchmark in Figure 5B uses separate subsets. Final admitted training-set sizes were not recorded in the saved histories.

| Dataset | Source | N | Role |
|---|---|---|---|
| Gold-standard positives | Li et al. 2026 | 40 | Source positive set |
| Gold-standard negatives | Li et al. 2026 | 14 | Source negative set |
| LOO-CV subset | Score-matched gold standard | 31 | 23 positives / 8 negatives |
| Primary variant table | Li et al. 2026 | 2,404 | Feature matrix |
| GENCODE v44 junctions | EBI/GENCODE | 252,835 | Configured extraction limit |
| gnomAD v4.1 benign | Broad Institute | ≤500 | Training candidate limit |
| MFASS | Chong et al. 2019 | ≤400 | Up to 200 per class |
| BRCA1 SGE | Findlay et al. 2018 | 200 | Cross-dataset eval |
| MaPSy | Soemedi et al. 2017 | 199 | Cross-dataset eval |
| VexSeq | Adamson et al. 2018 | 195 | Cross-dataset eval |
| SPiP | Leman et al. 2022 | 200 | Cross-dataset eval |
| GTEx v8 | GTEx Consortium | 54 tissues | Tissue expression |

**Table S6:** In-sample calibration metrics. Posterior means come from a Bayesian model fitted and evaluated on the same 31 variants. Calibration uses ten bins; only the two nonempty bins are listed. Confidence is the mean predicted disruption probability, and accuracy is the observed pathogenic fraction within each bin. These are in-sample rather than held-out LOO estimates.

| **Metric** | **Value** |  |  |
|---|---|---|---|
| Expected Calibration Error (ECE) | 0.040 |  |  |
| Maximum Calibration Error (MCE) | 0.040 |  |  |
| Brier Score | 0.002 |  |  |
| Number of bins | 10 |  |  |
| **Bin** | **Count** | **Confidence** | **Accuracy** |
| Bin 1 | 8 | 0.038 | 0.000 |
| Bin 10 | 23 | 0.960 | 1.000 |

**Table S7:** Fine-tuned diffusion-model performance. Results cover the full 54-variant gold standard and four external evaluation subsets. N_+_ and N*_−_*denote pathogenic and benign variant counts, respectively. The full gold standard is not an independent test set. These diffusion-model evaluations are distinct from 31-variant Bayesian LOO-CV and the 203-variant BRCA1 fine-tuning test. AUROC and AUPRC denote the areas under the receiver operating characteristic and precision–recall curves; Bal.Acc denotes balanced accuracy and MCC denotes the Matthews correlation coefficient.

| **Dataset** | **N** | **N<sub>+</sub>** | **N<sub>−</sub>** | **AUROC** | **AUPRC** | **Bal.Acc** | **MCC** |
|---|---|---|---|---|---|---|---|
| Gold standard | 54 | 40 | 14 | 0.600 | 0.859 | 0.679 | 0.319 |
| SPiP | 200 | 100 | 100 | 0.503 | 0.522 | 0.560 | 0.150 |
| MaPSy | 199 | 8 | 191 | 0.656 | 0.071 | 0.700 | 0.184 |
| VexSeq | 195 | 97 | 98 | 0.475 | 0.503 | 0.521 | 0.058 |
| BRCA1 SGE | 200 | 100 | 100 | 0.424 | 0.445 | 0.510 | 0.059 |

## SUPPLEMENTARY FIGURES

**Figure S1:**
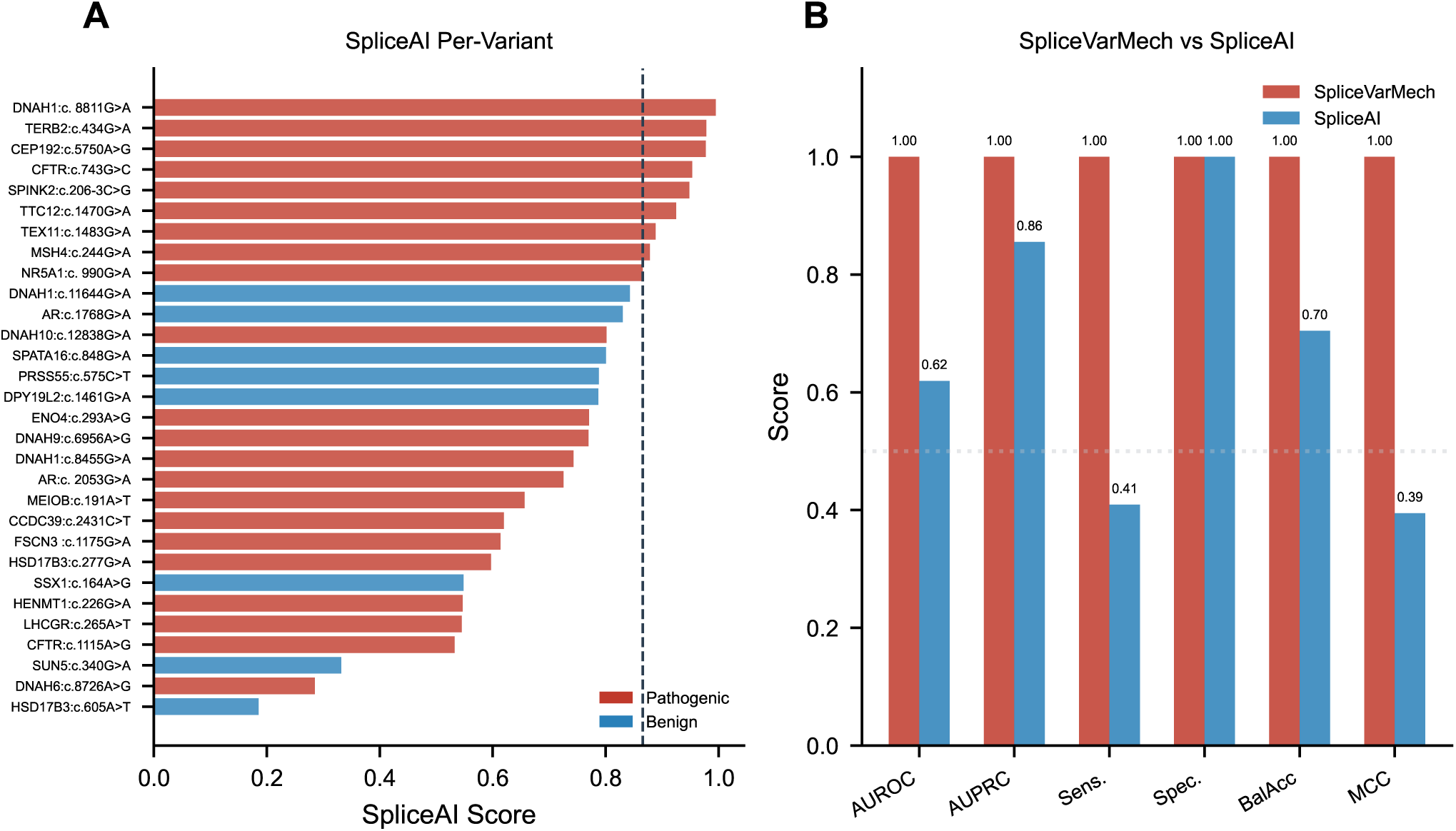
SpliceAI scores and comparative performance. (A) Each bar represents a SpliceAI score for one of the 30 scored variants, ordered by score. Red denotes pathogenic variants and blue denotes benign variants. The dashed line marks the optimal threshold of 0.866. (B) Summary metrics compare the 31 SpliceVarMech LOO predictions with SpliceAI results on its 30 scored variants. Red and blue bars identify SpliceVarMech and SpliceAI, respectively. Threshold-dependent metrics use the respective optimal thresholds of 0.410 and 0.866.

**Figure S2:**
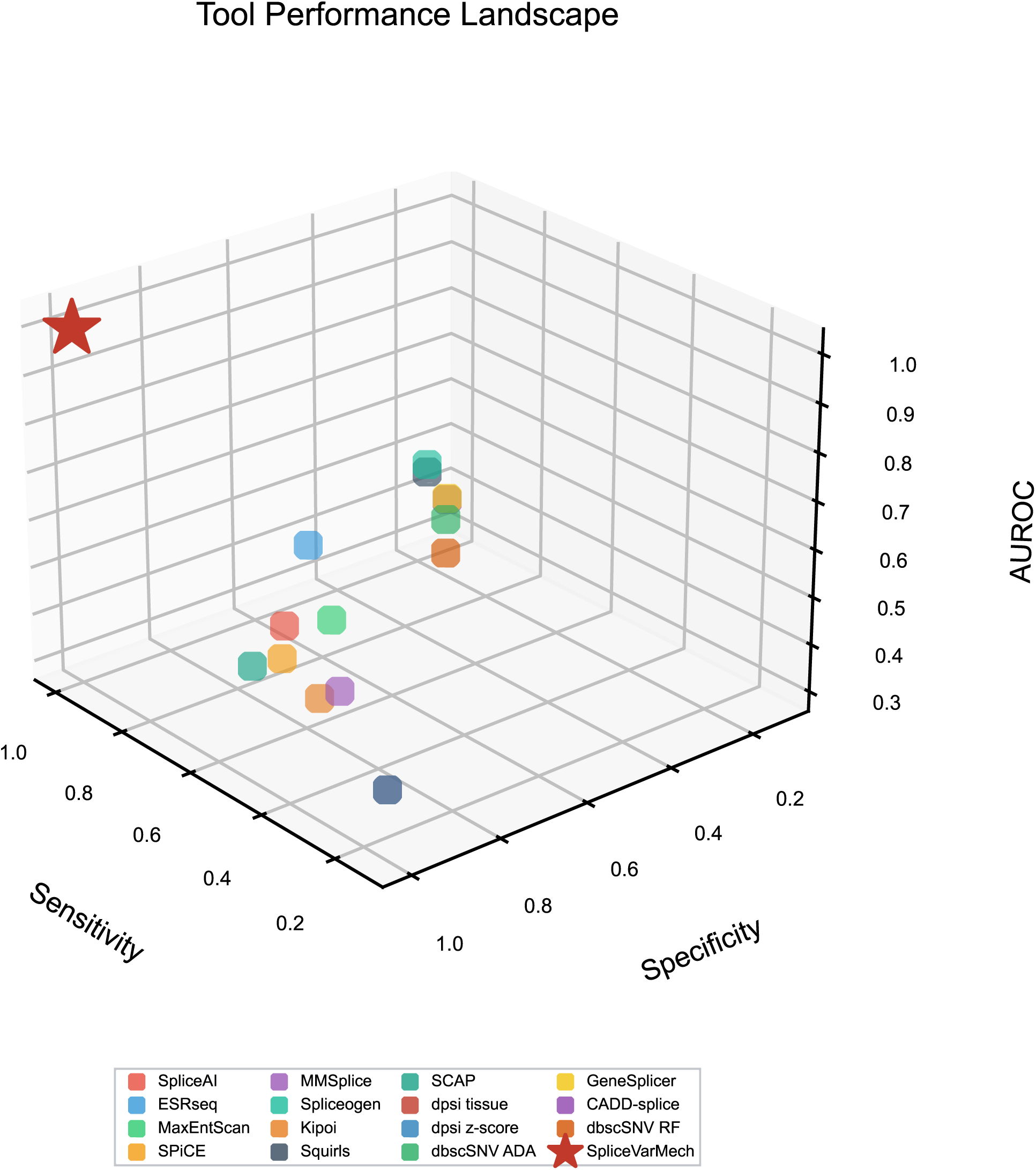
Baseline tool performance landscape. Each colored point represents an individual baseline tool with available sensitivity, specificity and AUROC on its scored subset of the 31-variant cohort. Sensitivity and specificity use each tool’s optimal threshold. The red star represents SpliceVarMech Bayesian LOO-CV, evaluated at threshold 0.410, with a value of 1.000 on each axis. Tools without estimable metrics are omitted.

**Figure S3:**
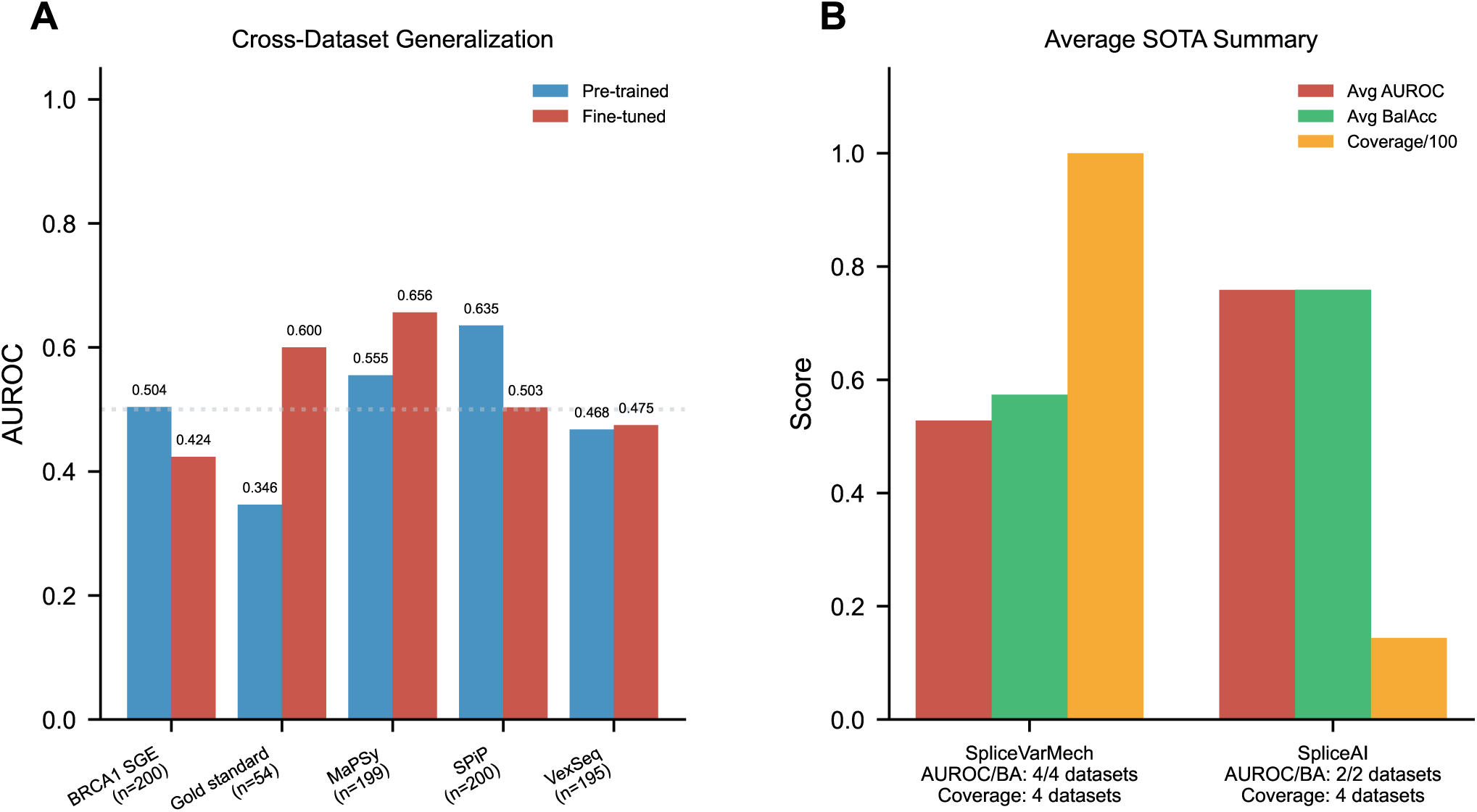
Model-stage comparison and benchmark averages. (A) Bars compare pre-trained and fine-tuned diffusion-model AUROC on the full gold standard and four external subsets. Blue denotes the pre-trained model and red denotes the fine-tuned model. Sample counts appear below the dataset names; these are the subsets summarized in Table S7. (B) Bars show unweighted means from the separate benchmark in Figure 5B. AUROC and balanced accuracy average four datasets for SpliceVarMech and the two datasets with available metrics for SpliceAI. Coverage averages all four datasets, treating unavailable SpliceAI coverage as zero, and is divided by 100 for display. Metric averages therefore use different numbers of datasets for the two methods.

## Notes

https://github.com/Moustafazn/AI-VUS-Mechanism

### Competing Interest Statement

The authors have declared no competing interest.

